# The influence of sex, hormone status, and growth rate on dietary isotope fractionation: a case study with laboratory rats

**DOI:** 10.1101/2021.03.25.436995

**Authors:** Sora L. Kim, Caroline Jones, Jessica Santollo

## Abstract

Stable isotope analysis is increasingly used to discern ecological differences within and among species, especially those difficult to observe. In applied ecological studies, variations in stable isotope composition are often attributed to foraging behavior or trophic ecology rather than fractionation during metabolic processes. One physiological difference among individuals is gonadal hormone levels, which affects food intake, metabolism, and locomotor activity. However, it is unclear how these effects– isolated from ecology– affect metabolic dynamics and expression in stable isotope analysis. Here, we test the linkage between gonadal hormones and isotopic heterogeneity among individuals with captive rats. We found that sex and removal of gonadal hormones are factors either individually or interactive for δ^15^N and δ^13^C values in serum and blood as well as δ^15^N values of muscle and liver. Furthermore, the degree of fractionation in these tissues is related to growth rate. Gonadectomization also affected isotopic composition for liver δ^13^C values and kidney δ^15^N values, but fat δ^13^C values were affected only by sex. The pattern of differentiation between groups was most different for kidney δ^15^N and δ^13^C values, which also had the largest isotopic variability among groups. Overall, isotopic variation within one tissue for the four groups demonstrated up to 1‰ difference in δ^13^C and δ^15^N values suggesting that sex based within population variations should into consideration these potential physiological effects.

**What is already known:** There are isotopic differences between diet and various consumer tissues with variability among individuals, which is often characterized based on captive feeding studies. Effects of protein quality and quantity, variable growth rate based on ontogeny, and differences between sexes have been tested thus far in experiments. Furthermore, it is known that gonadal hormones affect energy regulation with difference in weight gain, which suggest differences in extent and rate of biochemical processes.

**What this study adds:** This study demonstrates effects of gonadal hormone status with metabolic biochemistry by using stable isotopes of carbon and nitrogen as dietary tracers. We find that gonadal hormone status accounts for some isotopic heterogeneity among individuals, which should be considered when interpreting stable isotope data from wild specimens. Rather than attributing isotopic differences to ecological factors, such as foraging preference, there may be underlying physiological processes that differentiate groups based on ontogeny and sex.

## INTRODUCTION

Differences in foraging ecology can be difficult to assess within and between populations based on traditional dietary analysis techniques (i.e., observation, stomach-gut content analysis, fecal analysis). An increasing number of ecological studies rely on stable isotope analysis, a biogeochemical technique that uses natural abundance stable isotopes to track energy flow within an ecosystem. One underlying assumption when comparing stable isotope data (δ^13^C, δ^15^N) is that biochemical processes during metabolism transform dietary nutrients in a uniform pattern with limited variability among individuals within a group. Therefore, statistically significant differences in stable isotope data from field studies are often attributed to foraging behavior or trophic ecology, rather than physiological differences in dietary nutrient transformation.

The transformations associated with trophic transfer are combined into a biological parameter referred to as the discrimination factor (Δ^13^C, Δ^15^N). This parameter is empirically estimated from captive feeding experiments where isotopic composition is compared between diet and consumer (i.e., Δ^13^C = δ^13^C_consumer_ - δ^13^C_diet_). Experiments have explored the effects of sex (Kurle et al. 2014), tissue type (see review in Newsome et al. 2010), physiological stress (Hobson et al. 1993; Kempster et al. 2007), growth rate (Reich et al. 2008; Carleton and Martinez del Rio 2010), protein quality and quantity (Robbins et al. 2010; McMahon et al. 2015), and biochemical form of nitrogenous waste (see reviews by Minagawa and Wada 1984, Vander Zanden and Rasmussen 2001, McCutchan et al. 2003) but diet is often the manipulated factor. One trait discussed, but not directly tested to date, is hormone level within and between sexes. Research in laboratory rodents demonstrates sex differences in physiological parameters that effect energy homeostasis (Asarian and Geary, 2013; Shi et al. 2009), which in turn may affect isotopic fractionation associated with biochemical reactions during metabolism.

Locomotor activity, basal metabolism, and thermogenesis modulate energy expenditure. It is well established that activity levels in female rodents are greater than in males (Gentry and Wade 1976b; Ogawa et al. 2003). Female rodents also have higher VO_2_ consumption and CO_2_ release than males (Rodriguez-Cuenca et al. 2002; Valle et al. 2005; Shi et al. 2007), suggesting that female rodents have a higher basal metabolic rate. Finally, brown adipose tissue thermogenic activity is greater in females, compared to males (Rodriguez-Cuenca et al, 2002; Valle et al. 2005). This increased energy expenditure, coupled with lower daily food intake in females compared to males, results in sex differences in body weight (Gentry and Wade, 1976a). These sex differences are mediated by circulating gonadal hormones, sex chromosomes, and complex interactions between these factors (Shi et al. 2009; Asarian and Geary, 2013; Chen et al. 2013). Given these differences in the controls of energy homeostasis, it is reasonable to hypothesize that sex affects dietary isotope fractionation and macronutrient routing. One study reported sex differences in macronutrient routing in rats (Kurle et al. 2014), but the mechanism underlying this sex difference is unclear. Understanding the extent of stable isotope differentiation attributed to physiological processes is critical for proper interpretation of sex differences observed in ecological field studies using stable isotope analysis.

We aim to further investigate sex differences in dietary isotope fractionation and macronutrient routing by determining if gonadal hormones mediate any sex differences. We examined the extent of isotopic fractionation in serum, blood, liver, and kidney samples of sham operated control vs. gonadectomized male and female rats. All individuals were fed a standard chow diet throughout the experiment because we were not interested in incorporation rates and wanted to test for any baseline sex differences. Although isotopic differentiations were small among groups, there were effects on most tissues, which related to growth as well as sex and surgery.

## METHODS

### Animals and Housing

Sixteen age matched male and female Sprague Dawley rats (Envigo Laboratories, Indianapolis, IN), approximately 55 days of age upon arrival, were used in all studies. Rats were pair housed in standard plastic tub cages with ad libitum access to food (Tekland 2018; Harlan Laboratories) and tap water. Body weight was measured daily. The temperature and humidity controlled colony room was maintained on a 12:12 Light:Dark cycle (lights on at 0700 h). All experimental protocols were approved by the Animal Care and Use Committee at the University of Kentucky, and the handling and care of individual animals was in accordance with the *National Institutes of Health Guide for the Care and Use of Laboratory Animals*.

### Surgery

One week after arrival into the facility, all animals underwent sham (n = 8) or gonadectomy (n = 8) surgeries using an intraabdominal approach as describe previously (Santollo et al. 2018). Briefly, rats were anesthetized with a mixture of ketamine (80 mg/kg im; Henry Schein, Dublin, OH) and xylazine (4.6 mg/kg im; Akorn Animal Health, Lake Forest, IL) and received an injection of carprofen (5 mg/kg sc; Norbrook Labs, Newry, N. Ireland) as an analgesic. A small skin incision was made and the abdomen wall was opened. The ovaries and testes were externalized and removed or were placed back into the abdomen (sham-operation). The muscle wall was sutured and the skin was closed with wound clips. All animals then received a bolus injection of isotonic saline (sc, 5ml) for hydration and antisedan (1 mg/kg sc; Zoetis, Kalamazzo, MI) to reverse the sedative. All animals received a second injection of carprofen 24 h after surgery. Although all individuals received surgery, we will refer to those that underwent the gonadectomy procedure as the ‘surgery’ group, which includes castrated (CAST) and ovariectomized (OVX) individuals.

### Blood and Tissue Collection

Animals were allowed a two-week recovery period after surgery. On Test Day 1 the food in the animal’s cages was changed out to ensure that all food consumed during the 30 day test period was from the same lot (mill date 11/7/2016) of Tekland 2018 (Table 1). Tail blood was collected during the first half of the light phase on Days 1 and 15. Animals were briefly anesthetized with isoflurane and a 25G needle was inserted into the tail to collect 0.5 ml of blood. Blood was allowed to clot for 1 h, spun at 4°C for 15 min at 12,000 rpm, the serum and blood cells were collected, and then stored at 20°C until processing. On Test Day 30 animals were briefly anesthetized with isoflurane and then decapitated. Trunk blood was collected and serum and blood cells were collected as described. Muscle, fat, kidney, and liver samples were dissected out, rapidly frozen on dry ice, and stored at 20°C until processing.

**Table 1:** Nutritional composition of Tekland Global 18% Protein Rodent Diet.

| **Macronutrients** |  | **Main Ingredients** |
| --- | --- | --- |
| Crude Protein | 18.6% | Ground wheat,<br>ground corn, wheat<br>middlings, dehulled<br>soybean meal, corn<br>gluten meal, soybean<br>oil, brewers dried<br>yeast, minerals,<br>amino acids, and<br>vitamins |
| Fat | 6.2% |  |
| Carbohydrate | 44.2% |  |
| Crude Fiber | 3.5% |  |
| Neutral Detergent<br>Fiber | 14.7% |  |
| Ash | 5.3% |  |
| *Total* | *92.5%* |  |
| **Vitamins and**<br>**Minerals** | 7.5% |  |

### Stable Isotope Analysis

All samples were freeze dried overnight (LabConco Corp., Kansas City, MO) before stable isotope analysis. Rat tissues were weighed to 0.7-1.1 mg, except fat, which was weighed to 0.5 mg. Rat chow was homogenized with a ball mill (SPEX SamplePrep, Metuchen, NJ) and weighed to 1.5 mg for nitrogen and 0.5 mg for carbon isotope analysis. Samples were weighed on a Sartorius microbalance (0.0001mg precision) into tin capsules (3×5 mm from Costech Supplies) and analyzed at the University of Kentucky Stable Isotope Facility with a Costech 4010 Elemental Analyzer coupled with a Conflo III to a Thermo Finnigan Delta Plus XP continuous flow, isotope ratio mass spectrometer. All runs (n=7) included reference materials of known isotopic value (acetanilide [n>12], dogfish muscle tissue [DORM-3, n>3], and chicken feather [CCHIX, n>3]) for normalization, mass linearity, and drift corrections. The standard deviation for both δ^15^N and δ^13^C values were <0.1 within each run.

### Data Analysis

Data are presented as mean ± SEM throughout. Daily body weights were analyzed by a 3 factor ANOVA (sex X surgery X day). Body weights at the start and end of the experiment and average daily body weight change were analyzed with a 2 factor ANOVA (sex X surgery). The δ^15^N and δ^13^C values for each tissue were analyzed with a 2 factor ANOVA (sex X surgery). *A priori* post hoc tests along with Tukey’s post hoc tests were used throughout to follow up any statistically significant main or interactive effects. In addition, linear regressions were conducted to determine the relationship between growth rate vs. δ^15^N and δ^13^C values in each of the tissue samples.

## RESULTS

### Body Weight Gain

As expected, body weight on the day of surgery was greater in males, compared to females (F(1,12) = 702.94, p < 0.001). There was no difference in body weights between surgery groups (F(1,12) = 0.16, p = n.s.), nor any interactions between sex and surgery groups (F(1,12) = 0.25, p = n.s). (Figure 1A). Across the 30 day test period, daily body weight was influenced by sex (F(1,12) = 250.28), surgery (F(1,12) = 15.12), day (F(29,348) = 112.36), and an interaction between sex and surgery (F(1,12) = 8.25), day and sex (F(29, 348) = 8.32), and day and surgery (F(29,348) = 4.7), all ps < 0.01 (Figure 1A). Post hoc tests revealed that body weight increased as a function of days after the experiment commencement, males weighted more than females, gonadectomized animals weighted more than intact animals, and while there was no difference in body weight between the two groups of male rats, the gonadectomized females weighted more than the intact females, p < 0.05.

**Figure 1.**
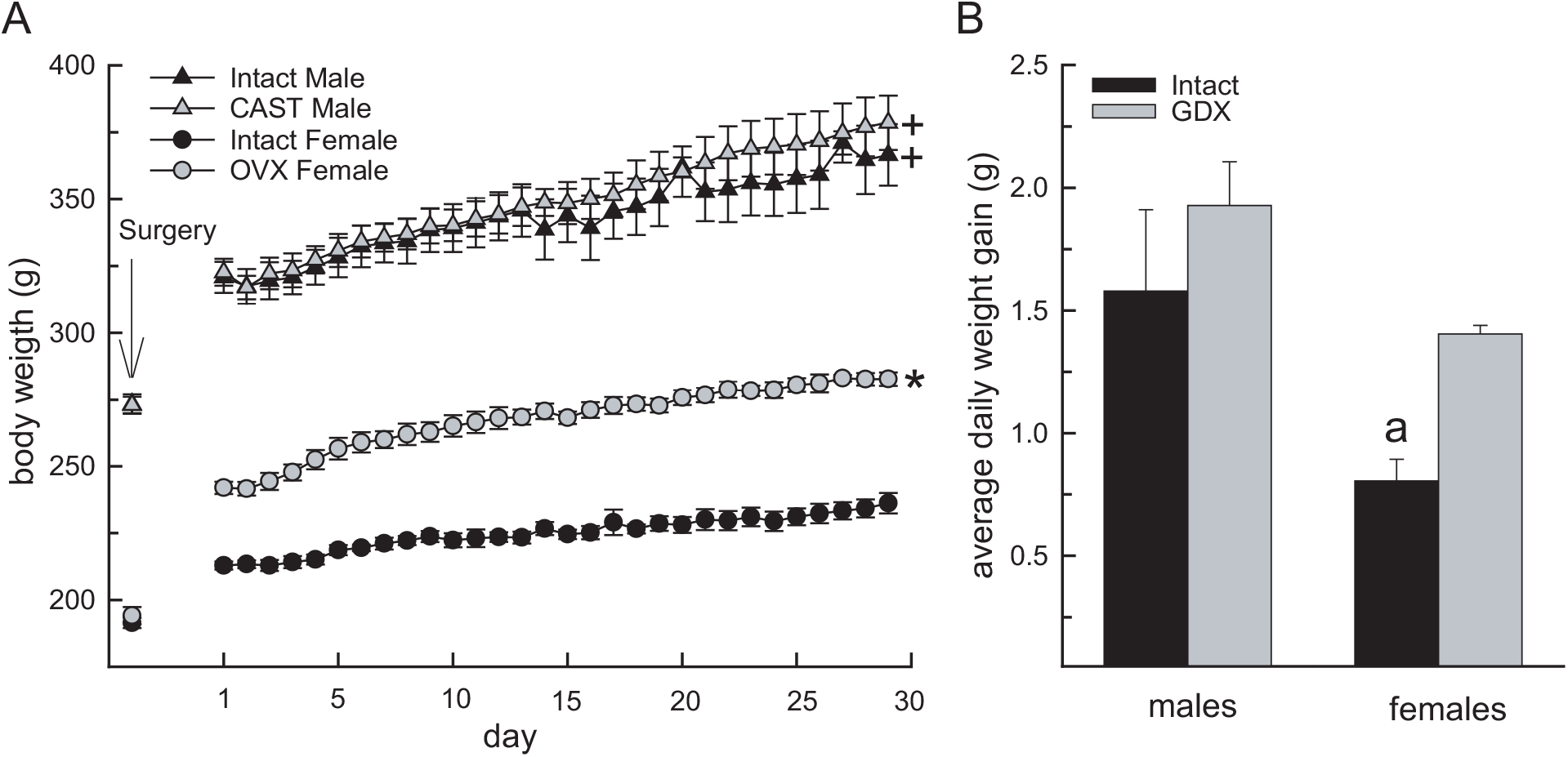
Body weight was influenced by sex and surgery. A) Daily body weight was greater in males, compared to females, but was not affected by gonadectomy. In females, daily body weight was greater in gonadectomized, compared to intact, rats. B) Average daily body weight gain was significantly lower in intact females compared to males and gonadectomized females. +Greater than females, p < 0.05. *Greater than intact females, p < 0.05. ^a^Less than all other groups, p < 0.05.

**Figure 2.**
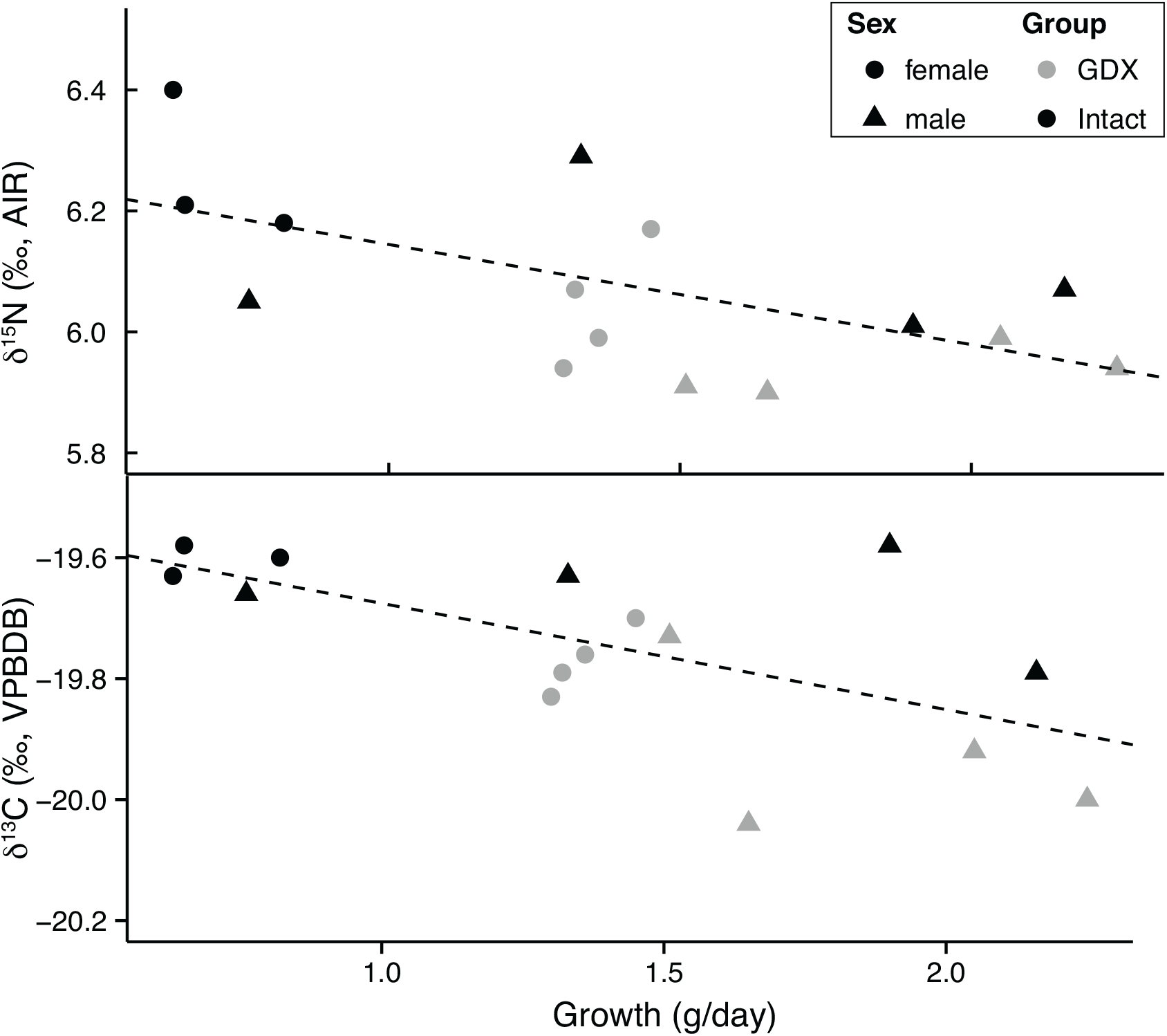
Serum δ^15^N and δ^13^C values are related to growth rate. There are differences in growth rate and isotope values with sex (□= female, ▴ =male) and gonadectomized individuals (GDX in grey).

Analysis of the average change in body weight revealed similar outcomes as the analysis of daily body weight. Daily body weight changes was influenced by main effects of sex (F(1,12) = 11.17, p < 0.01), and surgery (F(1,12) = 5.95, p < 0.05; Figure 1B). Post hoc tests revealed that daily body weight changes were greater in males than females and in gonadectomized animals compared to intact animals, p < 0.05. Finally, a prior post hoc testing revealed that daily body weight gain was significantly greater in both groups of males and the gonadectomized females compared to the intact females, p < 0.05.

### Stable Isotope Analysis

Serum, blood, kidney, liver, muscle, and fat samples were collected from this 30 day experiment; some tissues have longer incorporation rates than this period (residence time from Kurle [2009] is noted in Table 2), but we report all isotopic results. The isotope values for serum (days 0,15, and 30), blood (days 0,15, and 30), liver, kidney, muscle, and fat are reported in Table 2 and discrimination factors based on tissues at day 30 are reported in Table 3. The mean δ^15^N values for serum ranged as follows for days 0, 15, and 30: 5.6-6.1‰, 6.0-6.4‰, and 5.9-6.3‰. There was a correlation between growth rate and tissue (Table 2) as well as an interaction effect for time X sex X surgery (F(2,12) = 10.6, p < 0.01) where day 0 had lower δ^15^N values as well as a main effect of time (F(2,12) = 51.7, p < 0.0001). The mean δ^13^C values for serum ranged as follows for days 0, 15, and 30: -20.2--19.9‰, -20.1--19.8‰, and -19.9--19.6‰. For δ^13^C values of serum, there was also a correlation between growth rate and tissue (Table 2), but main effects were for time (F(2,12) = 27.3, p < 0.0001), sex (F(1,6) = 7.7, p < 0.0001), and surgery (F(1,6) = 13.9, p < 0.01) independently. The mean δ^15^N values for blood ranged as follows for days 0, 15, and 30: 3.9-4.3‰, 4.2-4.5‰, and 4.2-4.6‰. Blood δ^15^N values also had a correlation between growth rate and tissue (Table 2), interaction effects for time X surgery (F(2,20) = 13.5, p < 0.0001) with all time points difference from one another and sex X surgery (F(1,10) = 7.6, p < 0.05) with females greater than males, and main effects of time (F(2,20) = 108.4, p < 0.0001) and sex (F(1,10) = 11.5, p < 0.05). The mean δ^13^C values for blood ranged as follows for days 0, 15, and 30: -20.2--20.0‰, -20.1--19.9‰, and -20.0--19.7‰. Again, there was a correlation between growth rate and tissue (Table 2) and similar to serum δ^13^C results, there was main effects for time (F(2,20) = 138, p < 0.0001), sex (F(1,10) = 58, p < 0.0001), and surgery (F(1,10) = 11, p < 0.01) independently.

**Table 2.** Isotope values for various tissues sampled from rats in this experiment. Serum and blood were sampled on days 0, 15, and 30 as described in methods. All other tissues were sampled at the experiment’s termination. Residence time (1/λ) ranges are from nonlinear model results in Table 5 of Kurle (2009). Statistical results related to growth and isotope composition are also detailed. (ND = no data, ^‡^data from Kurle (2009))

| Tissue | Group | $\delta^{15}\text{N}$ | $\delta^{15}\text{N } 1\sigma$ | $\delta^{13}\text{C}$ | $\delta^{13}\text{C } 1\sigma$ | Statistical results<br>Growth | Statistical results<br>Stable isotope values |
| --- | --- | --- | --- | --- | --- | --- | --- |
| serum,<br>day 1 | OVX | 5.6 | 0.2 | -20.2 | 0.2 | $1/\lambda = 9\text{-}12^{\ddagger}$<br>N: $p = 0.01$, $r^2 = 0.58$<br>C: $p = 0.02$, $r^2 = 0.52$ | N: $p = \text{n.s.}$<br>C: $p = \text{n.s.}$ |
|  | Intact F | 6.1 | 0.1 | -19.9 | 0.1 |  |  |
|  | CAST | 5.7 | 0.1 | -20.2 | 0.1 |  |  |
|  | Intact M | 5.6 | 0.5 | -20.2 | 0.0 |  |  |
| serum,<br>day 15 | OVX | 6.1 | 0.1 | -20.0 | 0.1 | $1/\lambda = 9\text{-}12^{\ddagger}$<br>N: $p = 0.01$, $r^2 = 0.38$<br>C: $p = 0.005$, $r^2 = 0.44$ | N: **surgery** ($F(1,12) = 15.8$, $p < 0.05$), **sex** ($F(1,12) = 6.1$, $p < 0.05$)<br>C: **surgery** ($F(1,12) = 36$, $p < 0.0001$), **sex** ($F(1,12) = 7$, $p < 0.05$) |
|  | Intact F | 6.4 | 0.1 | -19.8 | 0.0 |  |  |
|  | CAST | 6.0 | 0.0 | -20.1 | 0.1 |  |  |
|  | Intact M | 6.1 | 0.2 | -19.9 | 0.0 |  |  |
| serum,<br>day 30 | OVX | 6.0 | 0.1 | -19.8 | 0.1 | $1/\lambda = 9\text{-}12^{\ddagger}$<br>N: $p = 0.02$, $r^2 = 0.35$<br>C: $p = 0.01$, $r^2 = 0.40$ | N: **surgery** ($F(1,12) = 14$, $p < 0.01$), **sex** ($F(1,12) = 6.5$, $p < 0.05$)<br>C: **surgery** ($F(1,12) = 20$, $p < 0.05$), **sex** ($F(1,12) = 5.1$, $p < 0.001$) |
|  | Intact F | 6.3 | 0.1 | -19.6 | 0.0 |  |  |
|  | CAST | 5.9 | 0.0 | -19.9 | 0.1 |  |  |
|  | Intact M | 6.1 | 0.1 | -19.7 | 0.1 |  |  |
| blood,<br>day 1 | OVX | 4.2 | 0.1 | -20.1 | 0.0 | $1/\lambda = 41\text{-}44^{\ddagger}$<br>N: $p = 0.04$, $r^2 = 0.31$<br>C: $p = 0.009$, $r^2 = 0.44$ | N: **sex** ($F(1,12) = 12.6$, $p < 0.01$), **sex X surgery** ($F(1,12) = 4.8$, $p = 0.05$)<br>C: **surgery** ($F(1,12) = 7$, $p < 0.05$), **sex** ($F(1,12) = 60$, $p < 0.001$) |
|  | Intact F | 4.3 | 0.1 | -20.0 | 0.0 |  |  |
|  | CAST | 4.1 | 0.0 | -20.2 | 0.0 |  |  |
|  | Intact M | 3.9 | 0.1 | -20.2 | 0.0 |  |  |
| blood,<br>day 15 | OVX | 4.2 | 0.2 | -19.9 | 0.0 | $1/\lambda = 41\text{-}44^{\ddagger}$<br>N: $p = 0.01$, $r^2 = 0.39$<br>C: $p = 0.04$, $r^2 = 0.26$ | N: **surgery** ($F(1,12) = 5.8$, $p < 0.05$), **sex** ($F(1,12) = 4.5$, $p = 0.05$), **sex X surgery** ($F(1,12) = 4.2$, $p = 0.063$)<br>C: **surgery** ($F(1,12) = 6$, $p < 0.05$), **sex** ($F(1,12) = 19$, $p = 0.001$) |
|  | Intact F | 4.5 | 0.1 | -19.9 | 0.1 |  |  |
|  | CAST | 4.2 | 0.1 | -20.1 | 0.1 |  |  |
|  | Intact M | 4.2 | 0.1 | -20.0 | 0.0 |  |  |
| blood,<br>day 30 | OVX | 4.4 | 0.1 | -19.9 | 0.1 | $1/\lambda = 41\text{-}44^{\ddagger}$<br>N: $p = 0.005$, $r^2 = 0.43$<br>C: $p = 0.004$, $r^2 = 0.45$ | N: **sex X surgery** ($F(1,12) = 8.2$, $p = 0.01$), **sex** ($F(1,12) = 24.3$, $p < 0.001$)<br>C: **surgery** ($F(1,12) = 14$, $p < 0.05$), **sex** ($F(1,12) = 44$, $p < 0.0001$) |
|  | Intact F | 4.6 | 0.1 | -19.7 | 0.0 |  |  |
|  | CAST | 4.3 | 0.1 | -20.0 | 0.0 |  |  |
|  | Intact M | 4.2 | 0.1 | -19.9 | 0.0 |  |  |
| liver | OVX | 6.0 | 0.2 | -21.2 | 0.0 | $1/\lambda = 4\text{-}11^{\ddagger}$<br>N: $p = 0.012$, $r^2 = 0.37$<br>C: $p = \text{n.s.}$ | N: **surgery** ($F(1,12) = 21.2$, $p < 0.001$), **sex** ($F(1,12) = 12.7$, $p < 0.05$)<br>C: **surgery** ($F(1,12) = 42.8$, $p < 0.001$) |
|  | Intact F | 6.2 | 0.1 | -21.0 | 0.1 |  |  |
|  | CAST | 5.6 | 0.2 | -21.3 | 0.1 |  |  |
|  | Intact M | 6.0 | 0.1 | -20.9 | 0.1 |  |  |
| kidney | OVX | 4.0 | 0.3 | -22.2 | 0.8 | 1/$\lambda$ = 24-54<sup>‡</sup><br>N: p = n.s.<br>C: p = n.s | N: **surgery** (F(1,12) = 9.2, p < 0.01)<br>C: n.s. |
|  | Intact F | 5.0 | 0.0 | -21.0 | 0.3 |  |  |
|  | CAST | 4.4 | 0.1 | -21.2 | 0.2 |  |  |
|  | Intact M | 4.9 | 0.6 | -21.0 | 0.2 |  |  |
| muscle | OVX | 4.6 | 0.1 | -20.0 | 0.1 | 1/$\lambda$ = 46-65<sup>‡</sup><br>N: p = 0.005, r<sup>2</sup> = 0.45<br>C: p = n.s. | N: **surgery** (F(1,12) = 11.3, p < 0.01), **sex** (F(1,12) = 13.2, p < 0.01)<br>C: n.s. |
|  | Intact F | 4.8 | 0.2 | -19.8 | 0.1 |  |  |
|  | CAST | 4.4 | 0.0 | -20.0 | 0.2 |  |  |
|  | Intact M | 4.6 | 0.1 | -19.3 | 0.2 |  |  |
| fat | OVX | -- | -- | -27.1 | 0.2 | 1/$\lambda$ : ND<br>N: NA<br>C: p = 0.06, r<sup>2</sup> = 0.22 | N: NA<br>C: **sex** (F(1,12) = 9.2, p = 0.01) |
|  | Intact F | -- | -- | -27.3 | 0.2 |  |  |
|  | CAST | -- | -- | -27.5 | 0.2 |  |  |
|  | Intact M | -- | -- | -27.5 | 0.2 |  |  |

**Table 3.** Discrimination values estimated from rats in this study fed standard Tekland Global chow (as described in Table 1).

| tissue | group | $\Delta^{15}\text{N}$ | $\Delta^{15}\text{N } 1\sigma$ | $\Delta^{13}\text{C}$ | $\Delta^{13}\text{C } 1\sigma$ |
| --- | --- | --- | --- | --- | --- |
| serum,<br>day 30 | OVX | 3.7 | 0.2 | 3.0 | 0.3 |
|  | Intact F | 4.0 | 0.2 | 3.2 | 0.3 |
|  | CAST | 3.6 | 0.1 | 2.9 | 0.3 |
|  | Intact M | 3.8 | 0.2 | 3.1 | 0.3 |
| blood,<br>day 30 | OVX | 2.1 | 0.2 | 2.9 | 0.3 |
|  | Intact F | 2.3 | 0.2 | 3.1 | 0.3 |
|  | CAST | 2.0 | 0.2 | 2.8 | 0.3 |
|  | Intact M | 1.9 | 0.2 | 2.9 | 0.3 |
| liver | OVX | 3.7 | 0.2 | 1.6 | 0.3 |
|  | Intact F | 3.9 | 0.2 | 1.8 | 0.3 |
|  | CAST | 3.3 | 0.2 | 1.5 | 0.3 |
|  | Intact M | 3.7 | 0.2 | 1.9 | 0.3 |
| kidney | OVX | 1.7 | 0.3 | 0.6 | 0.9 |
|  | Intact F | 2.7 | 0.1 | 1.8 | 0.4 |
|  | CAST | 2.1 | 0.2 | 1.7 | 0.4 |
|  | Intact M | 2.6 | 0.6 | 1.9 | 0.4 |
| muscle | OVX | 2.3 | 0.2 | 2.8 | 0.3 |
|  | Intact F | 2.5 | 0.2 | 3.0 | 0.3 |
|  | CAST | 2.1 | 0.1 | 2.8 | 0.3 |
|  | Intact M | 2.3 | 0.2 | 3.5 | 0.3 |
| fat | OVX | -- | -- | -4.3 | 0.4 |
|  | Intact F | -- | -- | -4.5 | 0.4 |
|  | CAST | -- | -- | -4.7 | 0.4 |
|  | Intact M | -- | -- | -4.7 | 0.4 |

We also consider final experiment results for serum, blood, kidney, liver, muscle, and fat without a temporal component. If only serum and blood day 30 results are considered, surgery and sex are main effects for serum δ^15^N, serum δ^13^C, and blood δ^13^C values whereas blood δ^15^N values have an interaction effect of sex X surgery and main effect from sex (Table 2). For liver, the mean δ^15^N values range 5.6-6.2‰ with main effects from surgery and sex whereas mean δ^13^C values range -20.9 to--21.3‰ with a main effect of surgery (Table 2). For kidney, the mean δ^15^N values range 4.0-5.0‰ with main effects from surgery whereas mean δ^13^C values range - 22.2--21.0‰ with no significant differences for any effects (Table 2). For muscle, the mean δ^15^N values range 4.4-4.8‰ with main effects from surgery and mean δ^13^C values range -20.1--19.8‰ with no significant differences for any effects (Table 2). Similar to serum and blood results, liver and muscle δ^15^N values were correlated to growth rate (Table 2). Finally, there are no δ^15^N values for fat because it is dominated by carbon, but δ^13^C values range -27.15--27.1‰ with a main effect of sex (Table 2). Although isotopic variation among groups is small, the replicates and statistical differences with sex and surgery as well as correlations with growth rate provide insights to the contribution of hormone status to metabolic dynamics during dietary nutrient incorporation. Finally, discrimination factors (Δ^15^N and Δ ^13^C) of tissues collected on day 30 were estimated for each group based on δ^15^N values (2.3‰, 1□ □= 0.1) and δ^13^C values (- 22.8‰, 1□ □ □ □ □ □ □) of diet chow pellets (n = 12); the reported error is based on propagation of uncertainty with standard deviation.

## DISCUSSION

Isotopic fractionation of dietary nutrients differed based on sex-specific gonadal hormone status with significant effects on multiple tissues. The largest factor related to significant isotopic differences was growth rate as gonadal hormone status affects weight gain. Tissues with faster incorporation rates exhibited significant differences among treatment groups in both δ^13^C and δ^15^N values (serum, blood, and liver), whereas tissues with longer incorporation rates only had significant differences among treatment groups for δ^15^N values (kidney and muscle). The differences observed among treatment groups are small, but this study is the first to explore the effects gonadal hormone status has on isotopic variation within a species and population.

### Growth Rate

The effect of gonadal hormone status on growth rate is well documented and suggests biochemical processes related to catabolism and anabolism differ. A natural tracer to explore the processes and mechanisms relating gonadal hormone status and metabolism is stable isotope analysis. Organic carbon and nitrogen isotopes trace biochemical reactions primarily related to amino acid synthesis. The body weight data during this experiment followed the trend of past studies where males were heavier than females and intact females gained the least amount of weight (Wade and Gentry 1978a). This growth trend suggests intact females have the highest energy expenditure (i.e., highest resting metabolic rate) and likely higher rates of catabolism, which would increase fractionation processes and produce higher isotope values and discrimination factors for consumer tissues than intact males or gonadectomized individuals. Because all individuals within this study were fed the same diet, we will discuss differences in isotope composition rather than discrimination factors, except when comparing results among studies (i.e., Kurle et al. 2014; Table 3).

Based on our growth data, we expected δ^15^N and δ^13^C values to be greatest in intact females with little to no distinction between gonadectomized males (CAST), gonadectomized females (OVX), or intact males. However, food intake varies among these groups (intact females < OVX < CAST < intact males; Gentry and Wade 1976a), which also plays a confounding factor in metabolizing nutrients. The pattern of variation between groups differed among tissues, but intact females had the highest δ^15^N values for serum, blood, liver, kidney, and muscle as expected from growth rate. For serum and liver, δ^15^N values are lowest for CAST while blood δ^15^N values are lowest for intact males, which suggests that feeding rate may be a more substantial factor than growth or these tissues were not in steady state as they only approached residence time (1/λ □ □ □ □ □ □ □ □ □) as reported in a previous incorporation rate study (Kurle et al. 2009). The carbon isotope dynamics also show some patterns; intact females have the highest δ ^13^C values for serum and blood and CAST have the lowest δ^13^C values for serum, blood, and liver. Similar to serum and blood δ^13^C values, fat also demonstrated a significant effect from sex, but it was the only tissue with no effect from surgery. The tissue that most differed from the expected isotopic pattern was kidney where δ^□5^N□ and δ^□ □^□ □ □ □ □ □ □ □ were lowest for the OVX group and intact males and females were similar with higher isotope values. In addition, the isotope composition of kidney tissue had higher variation than other tissues. The residence time of kidney tissue previously reported is 24-54 days (Kurle et al. 2009); these incorporation rates suggest that the kidney tissue may not have reached steady state with diet through the duration of this experiment. However, this differentiation in kidney isotope composition and variation among groups suggest this organ’s function may influence the mechanism driving the overall pattern in this experiment. Overall, our results elucidate the occurrence of higher δ^15^N and δ^13^C values in gonadectomized female (OVX) tissues as well as differentiation between gonadectomized males (CAST) and intact males, which suggests that estrogen, progesterone, testosterone, and other gonadal hormones play a factor in growth and metabolic processes.

### Comparison with previous discrimination factors

A previous stable isotope feeding study with rats varied protein and sugar sources to determine tissue specific discrimination factors (Kurle et al. 2014). Similar to our results, Kurle et al. (2014) observed lower Δ^15^N values for male rats across four diets and attributed this result to their increased growth rate. Furthermore, the experimental design controlled for protein quantity and quality, but found that protein source (i.e., wheat gluten, fish meal, wheat gluten/fish meal, or wheat gluten/fish meal/cow casein/chicken egg) affected Δ^15^N values while differences in source δ^13^C values caused variation in Δ^13^C values among tissues. In contrast to Kurle et al. (2014), this study kept diet constant and manipulated metabolic processes via gonadal hormone status.

The discrimination factors reported in Kurle et al. (2014) varied 1-2‰ per tissue and among sexes, which is greater variability than we measured within our experiment. Although serum Δ^15^N, liver Δ^13^C, and kidney Δ^13^C were similar between the two studies, serum Δ^13^C, blood Δ^15^N, blood Δ^13^C, and liver Δ^15^N values were greater in this experiment than reported in Kurle et al. (2014). Explanations for these differences could be related to internal (i.e., genetic background, life stage, or gut microbiome) or external (i.e., protein quantity, protein quality, or sample preparation) characteristics.

Many processes and mechanisms related to isotope fractionation between diet and consumer tissue remain enigmatic and are bundled within discrimination factors. We assessed the influence of gonadal hormone status, but there are other components, which may attribute to differences in discrimination factors between our study and Kurle et al. (2014). For example, while Sprague Dawley rats were used in both studies, they were obtained from different vendors; rats in the present study came from Envigo while rats in the Kurle et al. (2014) study came from Charles River. The genetic variability of outbred stocks, such as the Sprague Dawley rat, is considered to be even greater between different vendors due to different breeding practices. Although rats from these sources have similar body weight at 6 weeks, there is a divergence at 10 weeks and Charles River Laboratory rats are 24% heavier by 24 weeks despite similar food intake rates (Brower et al. 2015). The similarity in food intake but faster growth rate and heavier body weight would yield predications of lower discrimination rates for individuals in Kurle et al. (2014), which is the case when comparing results for Kurle et al. (2014) wheat diet (most similar in composition to this study) vs. this study’s results. Furthermore, we note that differences were greatest in serum Δ^13^C and blood Δ^13^C values suggesting that growth rate impacts both nitrogen and carbon fractionation during metabolic processes. Growth rate also likely differed due to life stage as there were substantial differences in the rats’ ages during the experiments; this study was conducted over 30 days versus 276-278 days (Kurle et al. 2014). Furthermore, the age during tissue sampling differed dramatically between studies. Young adult rats were used in the present study compared to middle-aged rats sampled in the study by Kurle and colleagues (2014). Finally, studies have shown differences in gut microbiome between male and female rats (Jašarević et al. 2016) with effects from gonadal hormones (Davey et al. 2012; Markle et al. 2013). There are also microbiome differences with genomic relationships to carbohydrate and protein metabolism as well as implications for plant vs. animal derived protein sources (Wang et al. 2014) that would affect the incorporation of nutrients between the standard diet rats received in this study vs. the specially formulated diets rats received in Kurle et al. (2014). Physiological processes tied to genetic background, life stage, growth rate, and gut microbiome influence discrimination factors among individuals and this variation must be considered when interpreting stable isotope data from wild animals.

The differences in discrimination factors between this study and Kurle et al. (2014) could also be rooted in external factors related to experimental design. Differences in nitrogen discrimination factors are often related to protein, especially in herbivorous or omnivorous species. Although protein quantity met the nutritional needs of the rats, the Tekland chow fed to individuals in this study was 18.6% while diets in Kurle et al. (2014) had protein content of 23.5-25.2%. It should be noted that the discrimination factors we estimated for serum and blood in this study, where diet protein was wheat based (table 1), were most similar to the wheat/fish diet from Kurle et al. (2014); this similarity demonstrates the influence of protein type on discrimination factors. Finally, samples were prepared differently between the two studies; we chose not to lipid extract samples but Kurle et al. (2014) performed chemical extractions with petroleum ether. Lipid is primarily composed of carbon that is ^13^C-depleted and therefore has lower δ^13^C values (Post 2007). Often, lipid is perceived as a confounding factor since content can skew δ^13^C values. However, our C:N ratios (based on weight %) were 3.5-3.7 for serum, 3.9-4.3 for blood, and 3.2-3.3 for muscle; for serum and muscle these values are similar to pure protein (3.2-3.4) and do not indicate lipid effects. The large differences in serum and blood Δ^13^C values between our study and Kurle et al. (2014) could be related to lipid extraction techniques. However, differences in Δ^15^N values should not be affected by lipid extraction.

### Potential mechanism: the role of insulin

Regardless of sex, removal of the gonadal hormones, and in turn removal of the major source of sex hormones, influenced metabolic processes as demonstrated by our isotopic analysis. The varying δ^15^N and δ^13^C values among treatment groups suggest differences in energy metabolism, which is tightly modulated by insulin, a hormone that helps maintain glucose levels within in a narrow range. Insulin affects carbohydrate, lipid, and protein metabolism and promotes substrate storage in liver, fat, and muscle while stimulating protein breakdown and synthesis (Champe et al. 2008). Gonadal hormones promote insulin signaling in a sex specific manner. In males, testosterone has direct effects of both pancreatic islet function and insulin responsive tissues by enhancing insulin gene and receptor expression and insulin release (Morimoto et al. 2001; Muthausamy et al. 2011). Ovarian hormones in females act in a similar fashion as testosterone in males. Estradiol promotes an increase in insulin synthesis and potentiates glucose-induced insulin release (Nadal et al. 2009; 2011). The tight relationship between insulin levels, energy metabolism, and gonadal hormone status suggests that these components factor into individual differences in food consumption, growth rate, and hence isotopic variability within a population and could explain how gonadectomy had a similar effect on isotopic fractionation processes in both males and females. Future studies will be needed to test the hypothesis that sex specific hormone effects on insulin signaling mediate the surgery effects observed here. Although many open questions remain related to the link between physiological processes and discrimination factors, this study lays the foundation for coupling stable isotope and hormone analysis in future applied ecological studies that focus on wild populations (i.e., Fleming et al., *in press*).

## ACKNOWLEDGMENTS

We thank Callie Whorf, Richard Dadundo, and Dr. Justin van de Velde for technical assistance. This work was supported by funds from the University of Kentucky College of Arts and Sciences to JS and SLK. A Summer Research and Creativity Grant from the University of Kentucky Office of Undergraduate Research supported CJ. Funds from the University of California Merced School of Natural Sciences supported SLK.

## DATA AVAILABILITY

https://doi.org/10.6071/M35379

